# Phenotypic and Kinetic Changes of Myeloid Lineage Cells in Innate Response to Chikungunya Infection in Cynomolgus Macaques

**DOI:** 10.1101/2021.05.18.444706

**Authors:** Brandon J. Beddingfield, Chie Sugimoto, Marcelo J. Kuroda, Eryu Wang, Scott C. Weaver, Kasi E. Russell-Lodrigue, Stephanie Z. Killeen, Chad J. Roy

## Abstract

Chikungunya is an emerging threat, spreading across multiple continents. The immune response to infection can lead to protection and convalescence, or result in long term sequelae such as arthritis. Early innate immune events during acute infection have been characterized for some cell types, but more must be elucidated with respect to cellular responses of monocytes and other myeloid lineage cells. In addition to their roles in protection and inflammation resolution, monocytes and macrophages are sites for viral replication and reservoirs for the virus. They are also found in joints post-infection, possibly playing a role in long term CHIKV-induced pathology. We examined kinetic and phenotypic changes in myeloid lineage cells, including monocytes, in cynomolgus macaques early after infection with CHIKV. We found increased turnover of monocytes and mDCs early during infection, with an accompanying decreases in both cell types, as well as a simultaneous increase in pDC number. An increase in CD16 and CD14 was seen along with a decrease in monocyte HLA-DR expression within 3 days of infection, potentially indicating monocyte deactivation. A transient decrease in T cells, B cells and NK cells correlates with lymphocytopenia observed during human infections with CHIKV. CD4+ T cell turnover decreased in blood, indicating relocation of cells to effector sites. This data indicates CHIKV influences turnover rates and kinetics of myeloid lineage cells early during infection, and may prove useful in development of therapeutics and evaluation of infection-induced pathogenesis.

## Introduction

Chikungunya is a rapidly emerging arboviral disease, with outbreaks occurring commonly in India and Southeast Asia^1^, that has resulted in approximately 1.9 million cases worldwide^2^. Since at least 2013, autochthonous transmission has occurred in the Americas^3^, with over 992,000 cases reported, including 332,000 in the United States and Canada (https://www.paho.org/data/index.php/en/mnu-topics/chikv-en/550-chikv-weekly-en.html). Clinical infection leads to a rapid onset, self-limiting, febrile illness with highly debilitating arthritic sequelae sometimes lasting months or years after infection^4,5^, with mortality generally resulting from neuroinvasive manifestations or severe organ dysfunction^6,7^.

Immune response to early CHIKV infection has been characterized with respect to natural killer (NK) cells, known to mediate the innate response against viral infection^8^. Clonal expansion of NK populations was highly correlated with viral titer in endemically-infected human populations^9^. NK cells are not necessary for protection from CHIKV infection^10^, instead displaying markedly altered expression of innate immune modulators such as IFN-□ ^9,11^, and may instead play a role in development of chronic arthritis arising from infection^12,13^.

A functional Type I Interferon response, important in the activation of monocytes and macrophages^14^, is necessary for protective immunity to CHIKV infection^15,16^. Monocytes and macrophages provide antiviral activities during infection^17,18^, as well as immunoregulatory functions to resolve inflammation^19^. In addition to their protective role during infection, monocytes and macrophages are likely to play a role in joint damage in CHIKV-induced arthritis, appearing in affected tissue along with mononuclear lymphocytes^13,20^. CHIKV actively replicates in monocytes/macrophages^21^, and persists within the cells long-term, with macrophages being major viral reservoirs^22^.

Dendritic cells (DCs) are an important cell type for early response to CHIKV infection^13^, with abrogated DC responses associated with increased disease severity^23^. Myeloid DCs (mDCs) exhibit a lowered number and delayed response in aged Rhesus macaques, coinciding with higher viral titers during infection^24^. Plasmacytoid dendritic cells (pDCs) also control CHIKV infection through Type I Interferon responses post CHIKV-infected cell contact, though NF-κB responses are excluded, thus potentially avoiding damage associated with other innate inflammatory responses^25^. This points to DCs as important players in the early control of infection.

In this study, we investigated innate immune responses by myeloid lineage cells, which include monocytes, macrophages, and dendritic cells in acute CHIKV infection that respond before the onset of adaptive immunity. We examined the phenotypic patterns and kinetics of monocytes, mDCs and pDCs during early Chikungunya infection in cynomolgus macaques. The data presented here will expand on current knowledge surrounding the innate immune responses to early CHIKV infection, which contribute to both pathogen clearance and long-term pathological consequence arising from this disease.

## Materials and Methods

### Animals

Age-matched cynomolgus macaques (*Macaca fascicularis*) weighing 3-6 kg, free of simian immunodeficiency virus (SIV), simian type D retrovirus (SRV), simian T-lymphotropic virus (STLV), and alphavirus antibodies against western (WEEV), Venezuelan (VEEV), and eastern equine encephalitis virus (EEEV), Sindbis virus (SINV), Semliki Forest virus (SFV), and CHIKV (assayed by hemagglutination inhibition) were used. The study was approved by the Institutional Animal Care and Use Committee at Tulane University, and all animals were handled in accordance with guidance from the American Association for Accreditation of Laboratory Animal Care.

### Viral Challenge

Anesthetized macaques were challenged with a single subcutaneous inoculation in the upper deltoid with wild-type CHIKV-LR (5.0 log_10_ PFU in a volume of 100 μl). Blood was collected on days 1-3, 6, 9, 13, and 35 after challenge, when the experiment was terminated and necropsies were performed. Tissues were placed in 10% zinc-formalin for histopathological analysis: brain, lung, heart, spleen, liver, kidney, mesenteric lymph node, bronchial lymph node, jejunum, colon, testes, muscle, skin, bone marrow, eye, and knee (or finger) joint. Some tissues were also frozen for viral titration by plaque assay: axillary, bronchial, and inguinal lymph nodes.

### BrdU inoculation

BrdU was dissolved in PBS at 30 mg/mL and inoculated intravenously at 60 mg/kg body weight two days post-challenge. EDTA blood was collected at 1, 4, 7 and 11 days post BrdU administration.

### Flow cytometric analysis

EDTA blood was stained for flow cytometric analysis. The following monoclonal antibodies (mAbs) were used in this study: anti-BrdU-FITC (3D4, BD Biosciences, San Jose, CA), CD45-PE (MB4-6D6, Miltenyi Biotec, San Jose, CA), CD20-ECD (B9E9, Beckman Coulter, Indianapolis, IN), CD123-PerCP Cy5.5 (BD), HLA-DR-PE-Cy7 (L243 BD Biosciences), CD11c-APC (BD), CD3-Alexa Fluor 700 (SP34-2, BD Biosciences), CD16-APC-H7 (3G8, BD Biosciences), CD14-Pacific Blue (M5E2, BD Biosciences), CD8-AmCyan (SK1, BD Biosciences), CD4-PerCP-Cy5.5 (L200, BD Biosciences), NKG2a-APC (Z199, Beckman Coulter). Blood was first stained with surface antibodies for 20 min. After lysing red blood cells with 1× FACS lysing solution (BD Biosciences), cells were permeabilized with Cytofix/Cytoperm (BD Biosciences) for 20 min, Permwash buffer (BD Pharmingen) supplemented with 10% dimethyl sulfoxide for 10 min, and Cytofix/Cytoperm for 5 min. The cells were incubated with DNase for 1 h at 37 °C, incubated with anti-BrdU antibody for 20 min, and fixed with 1% paraformaldehyde (Electron Microscopy Systems). Cells were acquired with LSR II (BD Biosciences) and analyzed using FlowJo software (TreeStar Inc).

### Complete blood counts and absolute counts of each cell population

The complete blood counts were analyzed on an ADVIA 120 Hematology System (Bayer Diagnostics) The absolute counts of CD14+ monocytes, mDC, pDC, CD4+ T cells, CD8+ T cells, B cells, and NK cells in blood were calculated from CBC data and flow cytometric analysis for each cell population.

### Statistical analysis

The statistical analysis was performed using Graphpad Prism software (GraphPad, San Diego, CA). Analysis was performed on post-exposure data defined as significant by the established detection thresholds for fever intensity, fever (hyperthermia) hours, and hypothermia hours. Fever intensity, indicated as the maximum change in temperature, fever hours, and hypothermia hours among treatment groups, was compared using the Kruskal-Wallis test. When the results of Kruskal-Wallis test indicated a significant difference at the *P*<0.05 level between the groups examined, Tukey’s multiple comparison test was then used to identify groups that differed at the *P*<0.05 significance level.

## Results

### Viral kinetics and gating strategy

Following challenge, virus levels were examined each day via plaque assay. Blood levels of CHIKV fell rapidly after initial peaks of 5.0 log_10_ at day 2 post challenge. By day 3 post challenge, viremia was absent in five out of seven animals (Table 1).

**Table 1.** Viremia in cynomolgus macaques after challenge with CHIKV.

| Viremia (log<sub>10</sub> PFU/ml) |  |  |  |
| --- | --- | --- | --- |
| Animal no. | Day 1 | Day 2 | Day 3 |
| HV78 | 4.3 | 5.0 | <1.0 |
| HV82 | 3.6 | 5.0 | <1.0 |
| HV84 | 3.3 | 5.0 | <1.0 |
| IV10 | 4.6 | 5.0 | 3.3 |
| IV11 | 4.0 | 4.6 | <1.0 |
| IV16 | 2.8 | 4.3 | <1.0 |
| IT87 | 5.2 | 5.3 | 3.5 |

Cells were subdivided based on gating of cell markers. Monocytes, mDC and pDC were identified as CD45+CD3-CD20-CD8-CD14+ cells, CD45+HLA-DR+Lin (CD3, CD20, CD8, and CD14)-CD11c+ cells, and CD45+HLA-DR+Lin-CD123+ cells, respectively. CD14 and CD16 expression pattern was used for subdividing monocyte population (Figure 1B).

**Figure 1.**
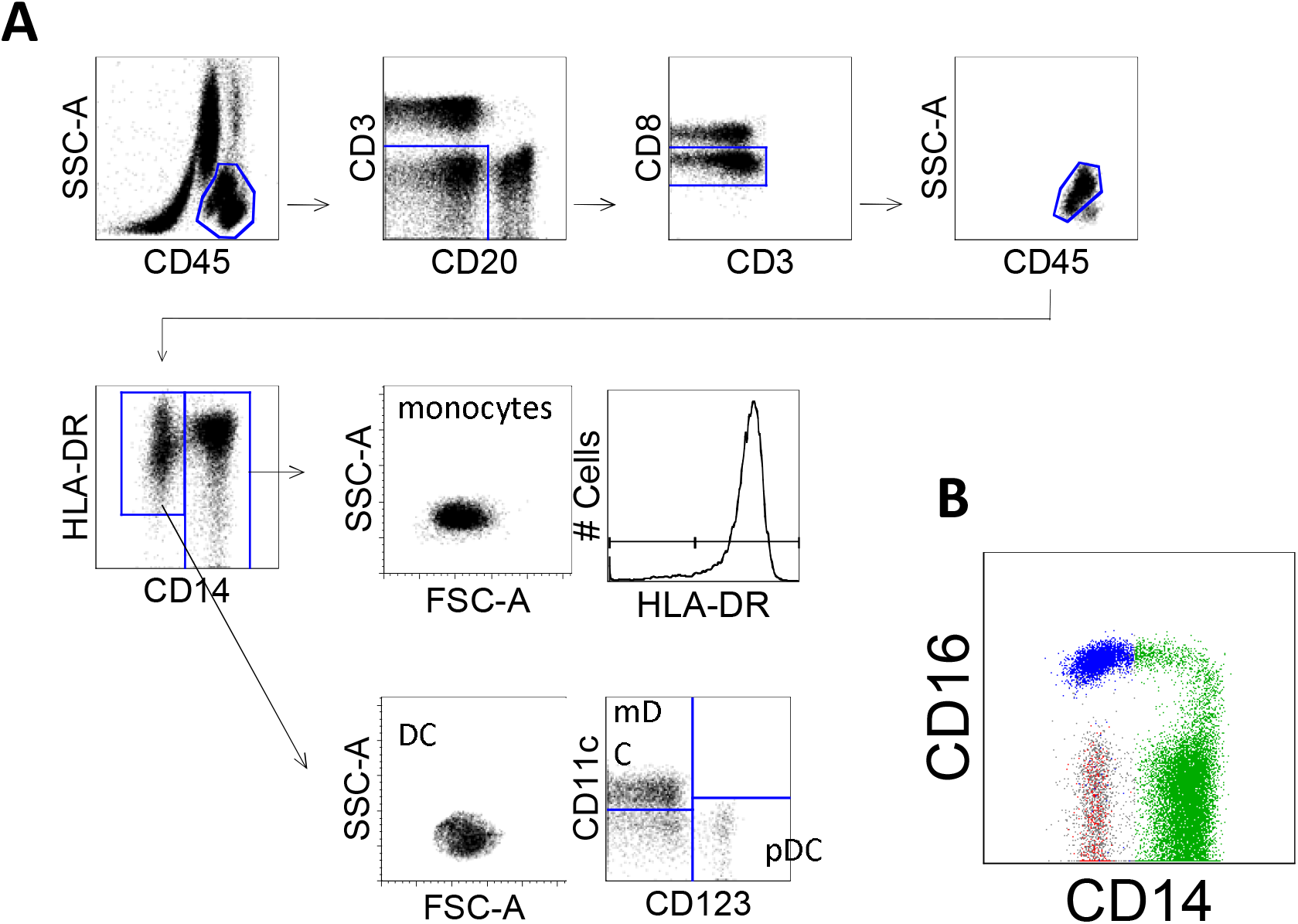
Gating strategy and phenotypes of myeloid lineage cells in peripheral blood from naive cynomolgus macaques. (A) Peripheral blood from naive cynomolgus macaques was analyzed by flow cytometry. Myeloid lineage cells which include monocytes and DC were gated as CD45^+^CD3^-^CD20^-^ CD8^-^. CD14 positive cells and CD14^-^HLA-DR^+^ were defined as monocytes and DC, respectively. DC were further divided into mDC and pDC by CD11c and CD123 expression. (B) Monocytes (green), mDC (blue)and pDC (red) were recognized as distinct populations on CD14 and CD16 expression.

### Turnover of myeloid lineage cells during acute infection

We examined kinetics of monocyte, mDC and pDC turnover by using *in vivo* BrdU labeling and flow cytometric analysis. BrdU was administered into CHIKV infected animals two days post challenge and blood was collected 1, 4, 7, and 11 days later. We compared the kinetics of the percentages of BrdU+ cells in monocytes, mDC and pDC between uninfected macaques and acutely CHIKV-infected macaques (Figure 2). Peripheral blood was collected at indicated time points (1 to 11 days) after BrdU administration and the appearance of BrdU+ cells was investigated in each cell population. Homogeneous kinetics of BrdU+ cells were observed in monocytes, mDC and pDC obtained from 8 uninfected macaques (Figure 2A). In contrast, the peaks of BrdU+ cells in all populations shifted to earlier time points in CHIKV-infected macaques, though this effect was only seen in two individuals for the monocyte population (Fig. 2A).

**Figure 2.**
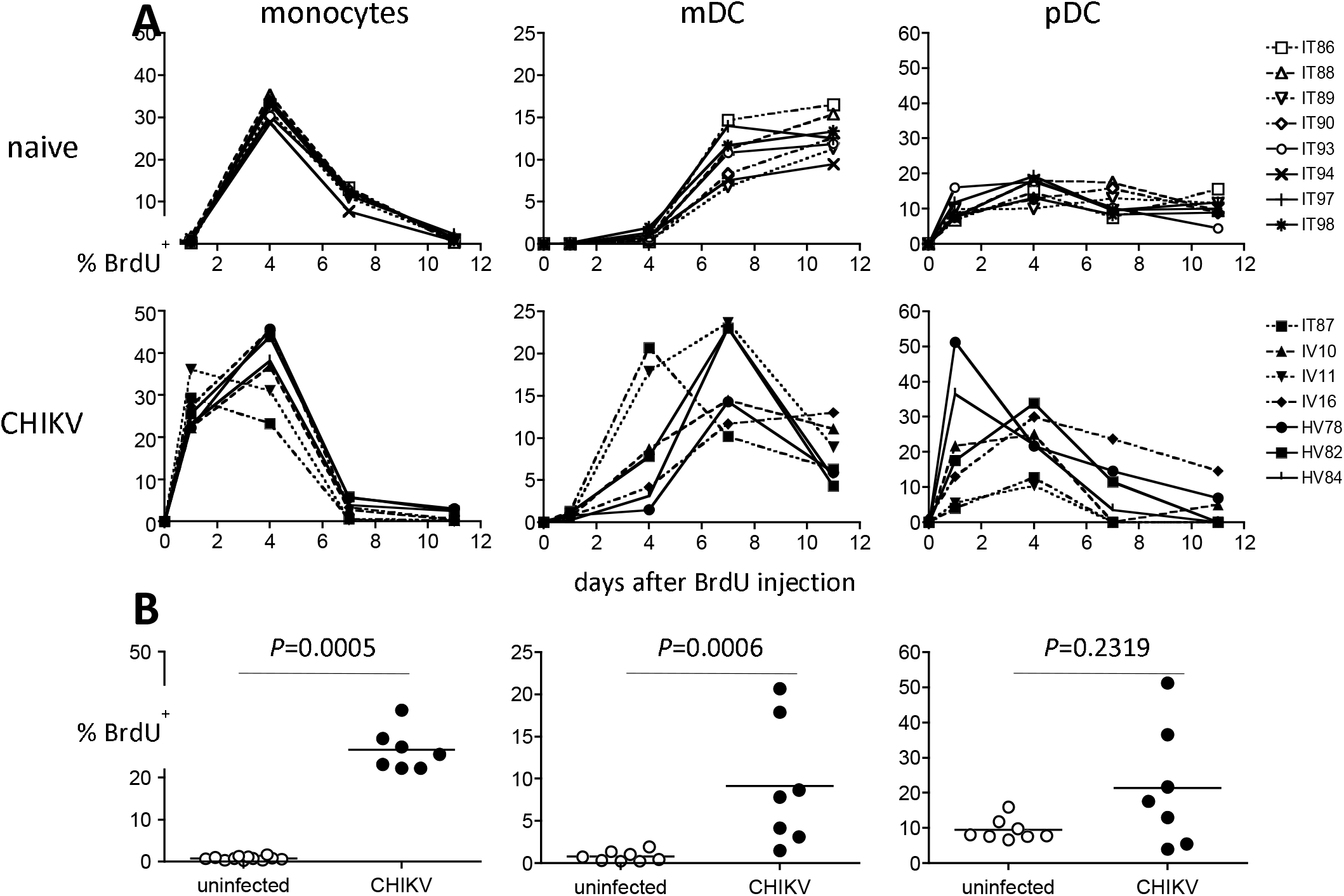
Kinetics of cell turnover of myeloid lineage cells in acute CHIKV infection. CHIKV-infected macaques were administrated BrdU at 2 days postinfection. Blood was collected serially after BrdU administration and stained for flow cytometric analysis. Monocytes, mDC, and pDC were identified as described in Figure 1. The percentages of BrdU positive cells were analyzed for each cell population. The statistical differences were analyzed by Mann-Whitney test with the data from 24h (for monocytes), 4 days (for mDC), and 24h (for pDC) after BrdU injection.

We analyzed the significance of increased cell turnover for monocytes, mDC, and pDC using the data from 24h, 4 days and 24h after BrdU injection, respectively. BrdU+ monocytes and mDCs in CHIKV-infected macaques were significantly higher than that from uninfected animals (Fig. 2B).

### Absolute cell counts of monocytes and DCs in acute CHIKV infection

Consistent with the Brd-based findings, monocyte counts decreased rapidly after challenge, but rebound with an increase 6 days after infection (Figure 3A). Dramatic changes were observed in mDC and pDC counts (Figure 3B and C, respectively), with mDCs largely disappearing from the blood immediately after infection, with blood pDC massively increasing at the same time.

**Figure 3.**
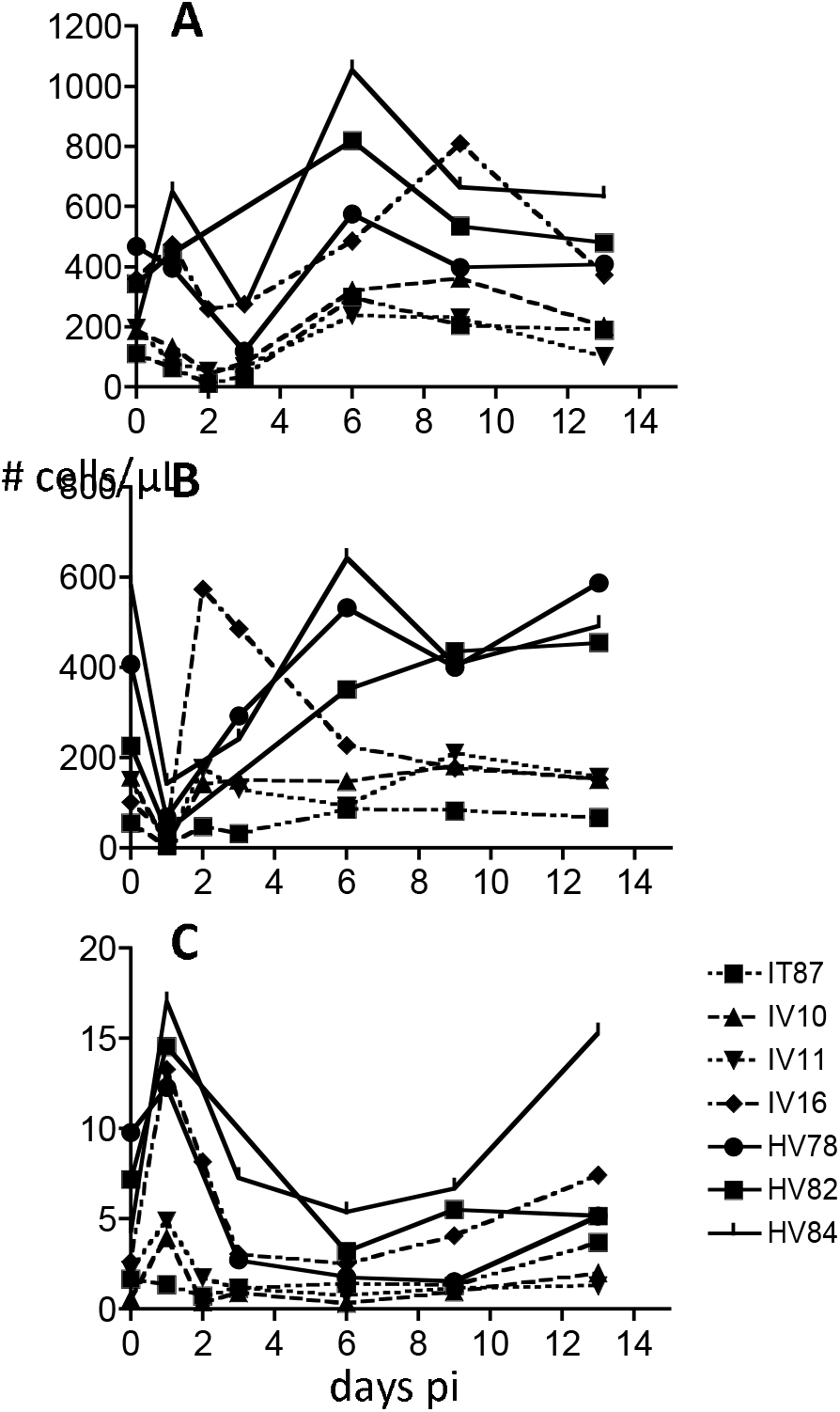
Absolute counts of monocytes (A), mDC (B), and pDC (C). Absolute counts of monocytes, mDC, and pDC were obtained from the calculation based on flow cytometric gating and complete blood counts (CBC).

### Phenotypic changes in monocytes from CHIKV-infected macaques

CD16 expression was up-regulated within 1 day after CHIKV infection and returned to the normal level roughly one week after infection (Figure 4A). Although we couldn’t obtain the blood samples at 2 days after infection in the first experimental group, 2 days post-CHIKV challenge was the peak of CD16 expression (Fig. 4B). CD14 expression was significantly higher 3 days post challenge, as compared to pre-challenge (Fig. 4C).

**Figure 4.**
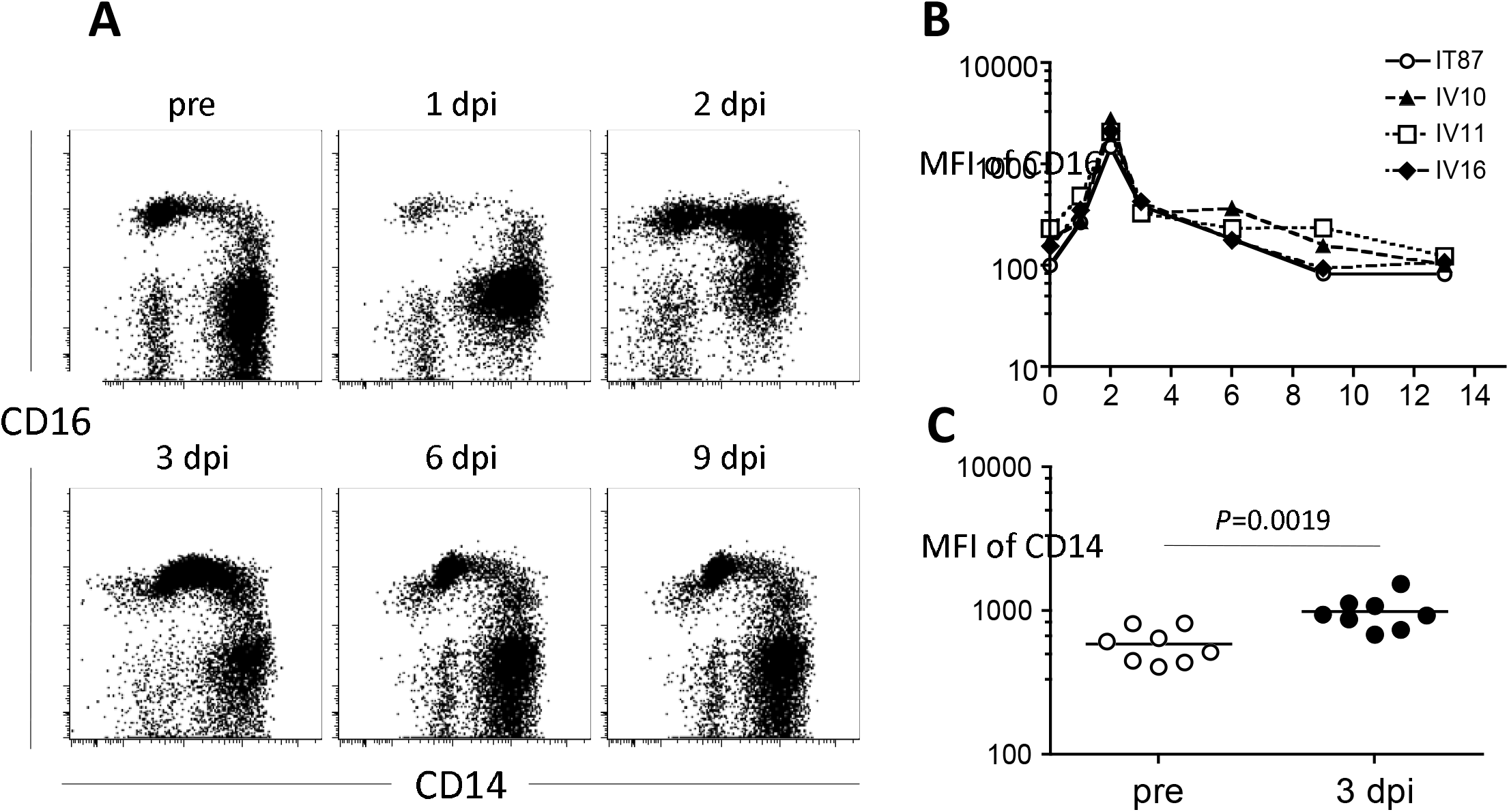
Altered CD14 and CD16 expression on myeloid lineage cell populations after CHIKV infection. The CD45^+^CD3^-^CD20^-^CD8^-^ population (as described in Figure 1) was examined for CD14 and CD16 expression by using serially collected blood after CHIKV infection. (A) Representative flow cytometric profiles from one of 7 CHIKV-infected macaques are shown. (B) Kinetic of mean fluorescence intensity of CD16 in R1 shown in (A) after CHIKV infection. (C) CD14 expression in R2 shown in (A) was compared between preinfection and 3 days after infection. The statistical difference was analyzed by Mann-Whitney test.

Furthermore, we determined that the HLA-DR expression on monocytes decreased in acute CHIKV infection (Figure 5). Although DC populations were HLA-DR-positive, the down-regulation of HLA-DR was not observed in either mDC or pDC (Fig. 5A). HLA-DR-negative or low monocytes appeared in 1 day after infection and returned to the uninfected level within 1 week after infection in most animals (Figure 4B). The majority of HLA-DR-down-regulated monocytes were CD16-negative (data not shown). There was no difference in the cell turnover between HLA-DR low/neg monocytes and HLA-DR hi monocytes (data not shown).

**Figure 5.**
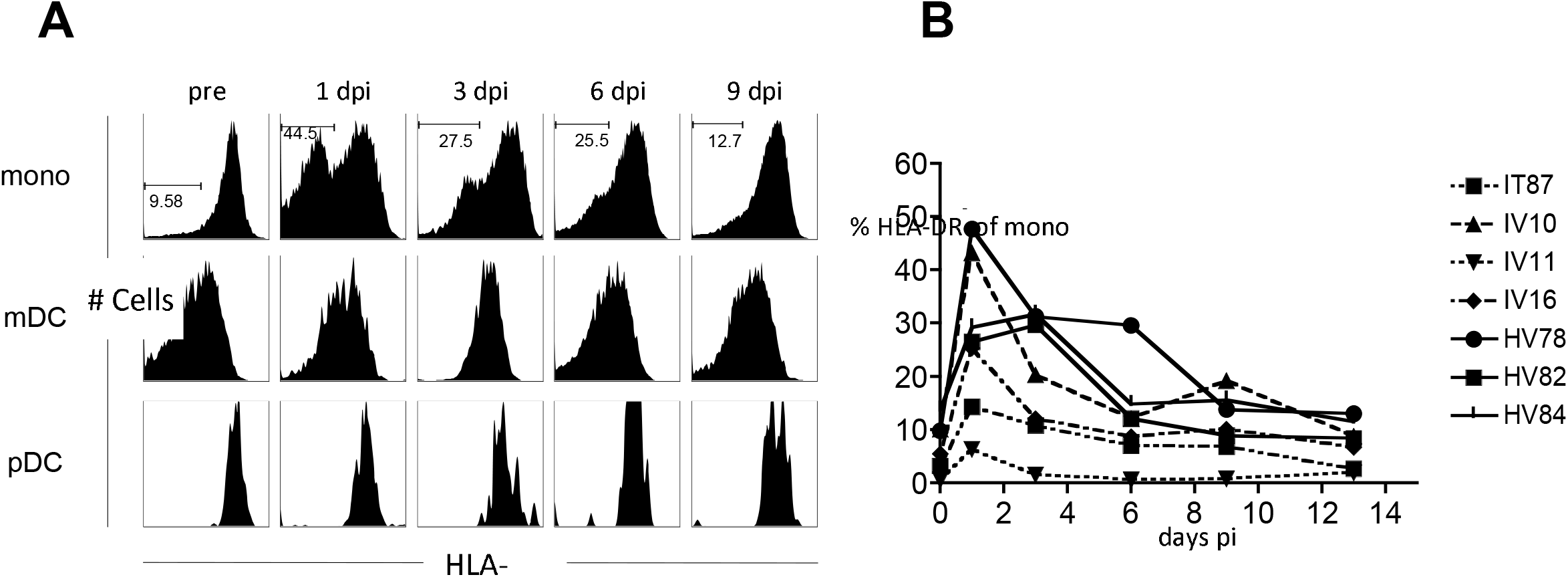
HLA-DR down-regulation in monocytes but not in mDC and pDC in acute CHIKV infection. Monocytes, mDC, and pDC were gated as Figure 1. The expression of HLA-DR was analyzed for each cell population. (A) Representative HLA-DR expression profiles on monocytes, mDC and pDC at the indicated days after infection are shown. (B) The percentages of HLA-DR low/negative of monocytes are plotted.

### Cell turnover and absolute counts of T cell, B cell and NK cell in acute CHIK infection

We examined the cell turnover and the absolute counts of the cell population related to adaptive immunity such as CD4+ T cells, CD8+ T cells, and B cells in blood. We also analyzed those for NK cells, since it has been reported that a phenotype change and a clonal expansion were observed in NK cells in acutely CHIKV-infected humans^9^. CD4+ T cells, CD8+ T cells, B cells, and NK cells temporarily decreased in a similar fashion after infection (Fig. 6).

**Figure 6.**
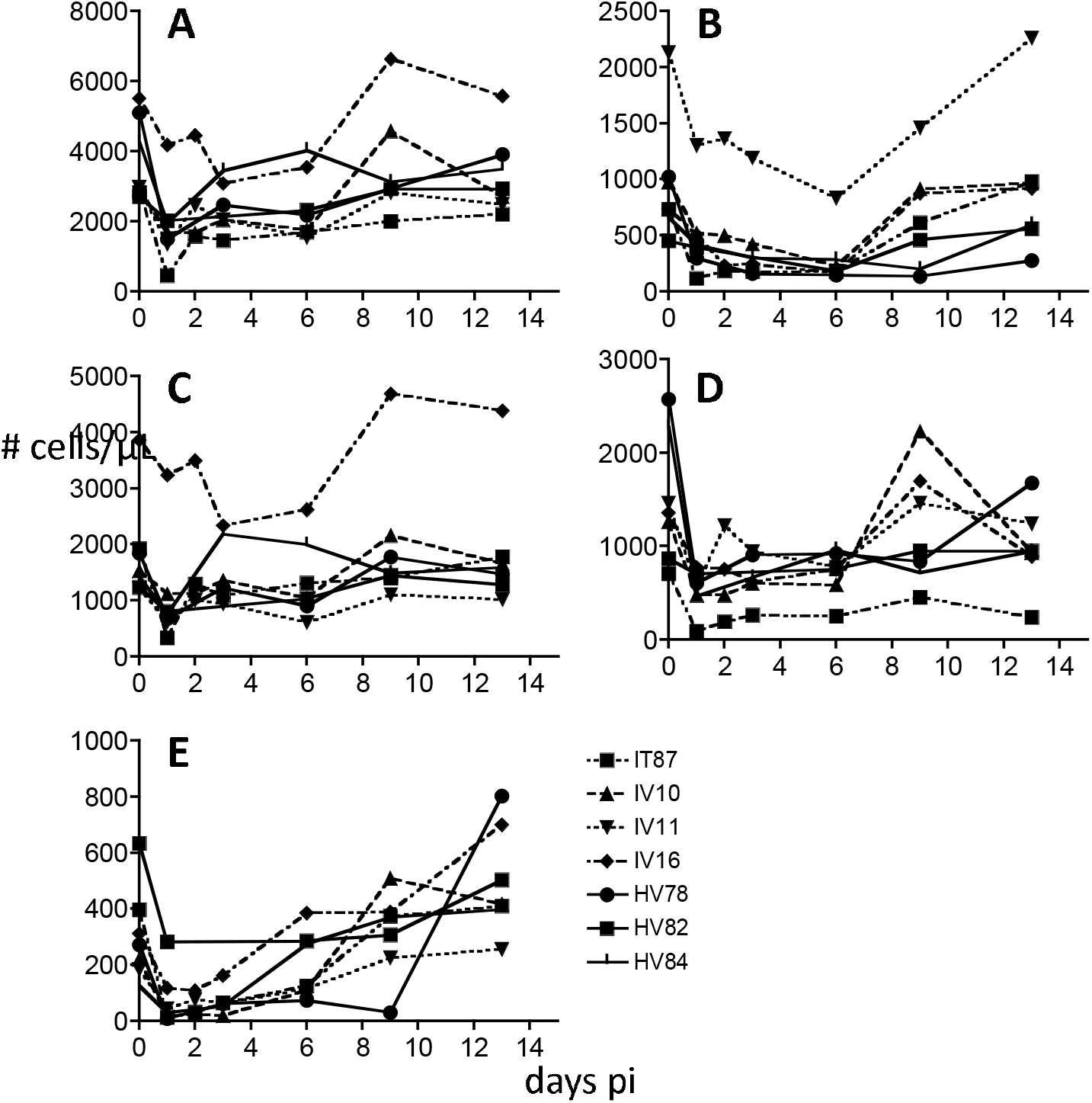
Absolute counts of CD3^+^ T cells (A), CD20^+^ B cells (B), CD4^+^ T cells (C), CD8^+^ T cells (D), and NK cells (E). Absolute count of each lymphocyte subset was obtained from the calculation based on flow cytometric gating (as described in Figure 6) and CBC.

Although the kinetics of the cell counts were similar, there were differences in the turnover among the cell populations. The cell turnover rates of CD4+ T cells were significantly decreased in acute infection (Fig. 7A), with CD8+ T cells also trending toward significant decreases (Fig. 7B). Cell turnover rates of B cells and NK cells were not significantly altered after infection (Figure 7C and D).

**Figure 7.**
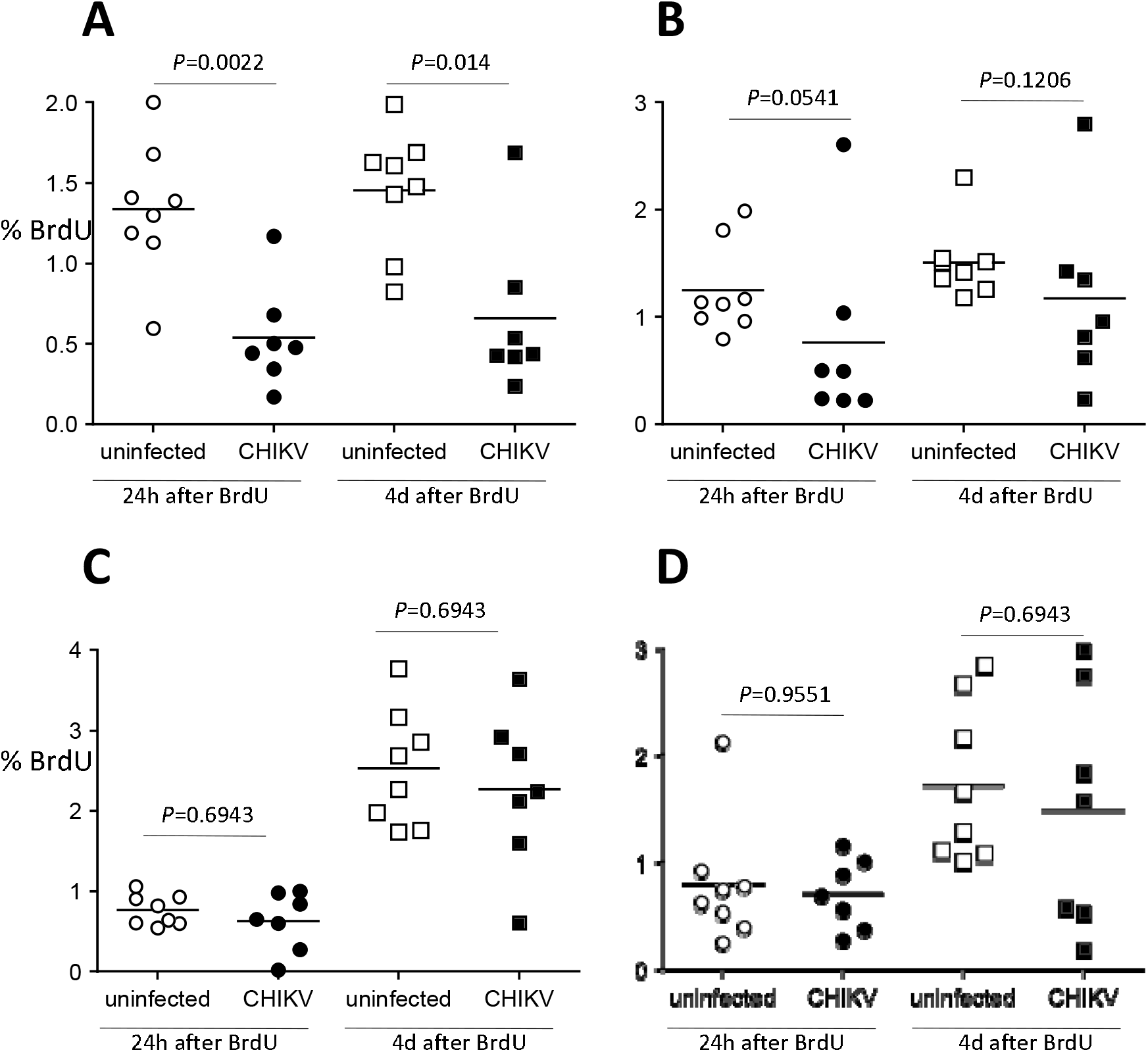
Lymphocyte turnover in acute CHIKV infection. CHIKV-infected macaques were administered BrdU at 2 days postinfection. Blood was collected serially after BrdU administration and stained with anti-CD3, CD4, CD8, CD20 and NKG2a, and BrdU antibodies for flow cytometric analysis. The data was compared to those form uninfected macaques. The cell turnover of CD4^+^ T cells (CD3^+^CD20^-^CD8^-^), CD8^+^ T cells (CD3^+^CD4^-^CD8^+^), CD20^+^ B cells (CD3^-^CD20^+^), and NK cells (CD3^-^ CD20^-^ CD8^+^NKG2a^+^) are shown in A, B, C, and D, respectively. The statistical differences between uninfected and CHIKV-infected groups were calculated by Mann-Whitney test.

## Discussion

The work presented here demonstrates myeloid lineage cell alteration in number, turnover and phenotype during infection with CHIKV. We demonstrated increased turnover of monocytes and mDCs early during CHIKV infection, which did not extend to pDC cells (Fig. 2). These changes tracked well with cell counts showing an early decrease of monocytes and mDCs, with an increase over time in all cell types, but an early increase of pDCs (Fig. 3). pDCs are known as type I IFN-producing cells that rapidly secrete massive amounts of type I IFN after activation via TLR7 and/or TLR9 by sensing ssRNA or CpG motifs in DNA derived from viral and bacterial genomes^26,27^. Therefore, extremely high CHIKV viral loads might result in an increased number of pDC, activation of pDC and induction of type I IFN by pDC. The early drop in circulating mDCs may be from increased tissue distribution, or an increase in apoptosis. Both are seen in SIV-infected macaques, along with a large drop in mDC number early during infection in one species of macaque, coinciding with more pathogenic infection; only a transient depletion is characterized during less severe disease^28^. We have found that the cell turnover in monocytes during SIV infection is highly correlated with progression to AIDS^29^, indicating monocyte/macrophage dynamics can play a large role in disease progression.

These myeloid lineage cell changes were accompanied by alterations in the phenotype of monocytes, with CD16 expression peaking at 2 days post-challenge, and CD14 expression peaking at day 3 (Fig. 4). CD16 is a marker of activation in monocytes, and corresponds to increases in pro-inflammatory cytokine secretion, which are seen in early CHIKV infection^30,31^. A considerable increase in the number of CD14+CD16+ monocytes had been described for a variety of systemic, infectious agents in humans, including hemolytic uremic syndrome^32^, bacterial sepsis^33^, HIV^34^, and experimental SIV infections in nonhuman primates^35^.

Monocytes expressing HLA-DR significantly dropped early during CHIKV infection, with a rebound seen by one week post challenge. This effect was not seen in dendritic cells (Fig. 5). Decreased HLA-DR expression is a hallmark of deactivated monocytes in patients with systemic inflammation, and this immune status is often referred to as compensatory anti-inflammatory response syndrome^36^. In *in vitro* studies, IL-10, an anti-inflammatory cytokine, induces HLA-DR down-regulation on monocytes^37^. Plasma IL-10 levels increased with increased induction of pro-inflammatory cytokines in acutely CHIKV-infected humans^38^.

We observed a temporary decrease in CD4+ and CD8+ T cells, as well as B cells and NK cells early during CHIKV infection (Fig. 6). This result corresponds to the lymphocytopenia observed in prior nonhuman primate models^22^, as well as in humans^39^. Turnover rates were altered during acute infection, with a decrease in rates of T cells, but no change in B cells or NK cells (Fig. 7). This suggests that proliferating CD4+ T cells and CD8+ T cells might preferentially relocate from the blood circulation to the effector sites.

These findings supplement previous work characterizing macrophages as virus reservoirs in cynomolgus macaques^22^. More detail regarding kinetics and phenotype of immune cells during infection with CHIKV can aid in the development of therapeutics, as well as provide more detail as to the relevance of non-human primate models to human infection.

## Conclusion

The early responses to chikungunya infection in the cynomolgus macaque are further examined here. Our findings suggest that this viral infection affects the production and turnover of myeloid lineage cells, especially monocytes and dendritic cells, as evidenced by their dramatic change in the absolute number and phenotypes early after infection. Future work should further elucidate how these phenotypic changes and turnover contribute to pathologic mechanisms of acute CHIKV infection, as well as its pathologic sequalae.

## Acknowledgements

This work was supported by a grant from the National Institute of Allergy and Infectious Disease (NIAID) through the Western Regional Center of Excellence for Biodefense and Emerging Infectious Disease Research, National Institutes of Health (NIH) grant U54 AIO57156. This work was also supported in part by NIH/NCRR grant number P51 RR000164.

